# Renal and Renal Sinus Fat Volumes as Quantified by Magnetic Resonance Imaging in Subjects with Prediabetes, Diabetes, and Normal Glucose Tolerance

**DOI:** 10.1101/620146

**Authors:** Mike Notohamiprodjo, Martin Goepfert, Susanne Will, Roberto Lorbeer, Fritz Schick, Wolfgang Rathmann, Petros Martirosian, Annette Peters, Katharina Müller-Peltzer, Andreas Helck, Susanne Rospleszcz, Fabian Bamberg

## Abstract

**Purpose:** The aim of this study was to assess the volume of the respective kidney compartments with particular interest in renal sinus fat as an early biomarker and to compare the distribution between individuals with normal glucose levels and individuals with prediabetes and diabetes.

**Material and Methods:** The sample comprised N = 366 participants who were either normoglycemic (N = 230), had prediabetes (N = 87) or diabetes (N =49), as determined by Oral Glucose Tolerance Test. Other covariates were obtained by standardized measurements and interviews. Whole-body MR measurements were performed on a 3 Tesla scanner. For assessment of the kidneys, a coronal T1w dual-echo Dixon and a coronal T2w single shot fast spin echo sequence were employed. Stepwise semi-automated segmentation of the kidneys on the Dixon-sequences was based on thresholding and geometric assumptions generating volumes for the kidneys and sinus fat. Inter- and intra-reader variability were determined on a subset of 40 subjects. Associations between glycemic status and renal volumes were evaluated by linear regression models, adjusted for other potential confounding variables. Furthermore, the association of renal volumes with visceral adipose tissue was assessed by linear regression models and Pearson’s correlation coefficient.

**Results:** Renal volume, renal sinus volume and renal sinus fat increased gradually from normoglycemic controls to individuals with prediabetes to individuals with diabetes (renal volume: 280.3±64.7 ml vs 303.7±67.4 ml vs 320.6±77.7ml, respectively, p < 0.001). After adjustment for age and sex, prediabetes and diabetes were significantly associated to increased renal volume, sinus volume (e.g. β_prediabetes_ = 10.1, 95% CI: [6.5, 13.7]; p<0.01, β_Diabetes_ = 11.86, 95% CI: [7.2, 16.5]; p<0.01) and sinus fat (e.g. β_prediabetes_ = 7.13, 95% CI: [4.5, 9.8]; p<0.001, β_Diabetes_ = 7.34, 95% CI: [4.0, 10.7]; p<0.001). Associations attenuated after adjustment for additional confounders were only significant for prediabetes and sinus volume (β =4.0 95% CI [0.4, 7.6]; p<0.05). Hypertension was significantly associated with increased sinus volume (β = 3.7, 95% CI: [0.4, 6.9; p<0.05]) and absolute sinus fat volume (β = 3.0, 95%CI: [0.7, 5.2]; p<0.05). GFR and all renal volumes were significantly associated as well as urine albumin levels and renal sinus volume (β = 1.6, 95% CI: [0.2, 3.0]; p<0.05). There was a highly significant association between VAT and the absolute sinus fat volume (β = 2.75, 95% CI: [2.3, 3.2]; p<0.01).

**Conclusion:** Renal volume and particularly renal sinus fat volume already increases significantly in prediabetic subjects. There is a significant association between VAT and renal sinus fat, suggesting that there are metabolic interactions between these perivascular fat compartments.

## Introduction

Parenchymal abnormalities of the kidneys are closely linked to the development and outcome of cardiovascular disease (1–6). Furthermore, chronic kidney disease itself is an independent risk factor for the development of coronary heart disease (1–3) and associated with adverse outcomes in existing cardiometabolic disease (4–6). The exact mechanisms that link obesity with insulin resistance, type 2 diabetes, hypertension, cardiovascular complications and renal diseases, are still not sufficiently clarified.

Perivascular fat, particularly intrahepatic fat and visceral adipose tissue (VAT) correlate with cardiometabolic risk factors, above and beyond standard anthropometric indices (7). Perivascular adipose tissue is in close contact with the adventitia of large, medium and small diameter arteries, possesses unique features differing from other fat depots and may also act independently of general obesity. Furthermore, there are locally acting fat depots such as peri- and epicardial fat, perivascular fat, and renal sinus fat (8). Renal sinus fat is thought to obstruct the blood and lymph outflow of the kidney and is associated with both cardiovascular risk factors such as hypertension, diabetes and chronic kidney disease (9) and might serve as an pathophysiological link and potential imaging biomarker.

Magnetic resonance imaging allows for excellent anatomical separation without the need of radiation or contrast agent administration. Multi-Echo sequences provide comprehensive image contrast, so that an in-phase, opposed-phase, fat only and water image is provided (10). This combination of contrasts allows for improved segmentation of the renal compartments. This methodology also forms the basis for absolute MR-quantification of renal sinus and intrarenal adipose tissue (11, 12). A semi-automated approach may yield a reliable method for high through-put quantification of the kidney compartments for large scale cohort studies.

Whole-body MRI has been implemented in population-based studies to detect early disease stages and imaging biomarkers indicative of increased risk of developing diseases in the future. Currently, a clinically well characterized subset of participants from the KORA cohort (Kooperative Gesundheitsforschung in der Region Augsburg) have undergone whole-body MRI for detection of atherosclerotic changes including cardiac function and morphometric and adipose tissue evaluation of abdominal organs such as the liver or the kidneys.

Thus, the aim of this study was to assess the volume of the respective kidney compartments with particular interest in renal sinus fat as an early metabolic biomarker and to compare the distribution between individuals with normal glucose levels and prediabetic and diabetic individuals.

## Material and Methods

### Study design

The study population consisted of a cross-sectional subsample of N = 400 whole-body MR participants from the population-based KORA FF4 cohort from the region of Augsburg, Germany. KORA FF4 (N = 2279, enrolled in 2013/2014) is the second follow-up of the original S4 survey (N = 4261, enrolled in 1999-2001, first follow-up: F4, N = 3080, enrolled in 2006-2008) (13). The setup of the MR substudy in KORA FF4 has been described previously (14). Eligible subjects were selected if they met the following inclusion criteria: willingness to undergo whole-body MRI; and qualification as being in the prediabetes, diabetes, or control group (see Covariate Assessment). Exclusion criteria were: age >72 years, subjects with validated/self-reported stroke, myocardial infarction, or revascularization; subjects with a cardiac pacemaker or implantable defibrillator, cerebral aneurysm clip, neural stimulator, any type of ear implant, an ocular foreign body, or any implanted device; pregnant or breast-feeding subjects; and subjects with claustrophobia, known allergy to gadolinium compounds, or serum creatinine level ≥ 1.3 mg/dl (14).

Data on VAT and intrahepatic fat was also derived from the same study. The study cohort has been analyzed in several other manuscripts (14–19). For detailed information please refer to the respective manuscripts.

This study was approved by the institutional review board of the Ludwig Maximilian’s University Munich (Germany) and written consent was obtained from each participant.

### MR imaging protocol

Whole-body MR measurements were performed on a 3 Tesla scanner (Magnetom Skyra, Siemens Healthcare, Erlangen, Germany). Detailed descriptions of technical and imaging protocols are listed elsewhere (14). For assessment of the kidneys, a coronal T1w dual-echo Dixon and a coronal T2w single shot fast spin echo (SS-FSE/HASTE) sequence were employed. Imaging parameters dual-echo Dixon: 256 x 256, field of view (FOV): 488 x 716 mm, echo time (TE) 1.26 ms and 2.49 ms, repetition time (TR): 4.06 ms, partition segments: 1.7 mm, flip angle: 9°. Image parameter T2 Haste: matrix: 320 x 200, field of view (FOV): 296 x 380 mm, echo time (TE) 91 ms, repetition time (TR): 1000 ms, partition segments: 5 mm, flip angle: 131°.

### Kidney Segmentation

The semi-automated image segmentation was performed using Matlab (Version R2011b, The MathWorks, Natick, USA) **(Fig 1)** and is based on the algorithm described in (20). In short, the kidneys were segmented from the surrounding tissues by thresholding the Dixon-T1 water-only images with a subsequent refinement step using prior knowledge about the kidney shape and location. In a second step renal parenchyma, renal sinus and sinus fat were determined by thresholding the maximum pixel’s intensity in the slice. The separation of the kidney from the spleen and gastrointestinal tract was refined using active contours generating a whole kidney mask. Within the generated entire kidney mask, the kidney, renal sinus and pelvis were subsequently separated. This separation algorithm utilizes assumptions of the renal anatomical structure, e.g., that the renal cortex surrounds parts of the medulla. The renal sinus was segmented using pixels with lower signal intensity than renal parenchyma tissue in water-only T1w-images. The fat-only pixels were identified through their position and separated from the pelvis mask. Afterwards, the union pelvis mask was subtracted from the kidney mask to separat kidney, renal sinus and pelvis. The resulting masks of the respective compartments were inspected by one reader (M.G.) and manually corrected by eliminating voxels mistakenly considered as renal parenchyma, mostly from the liver or spleen. The final volumes of the entire kidneys, renal cortex, medulla, and pelvis were then calculated by voxel summation. A subset of 33 study participants was also evaluated by a second reader (S.W.) for assessment of inter-reader variability.

**Figure 1.**
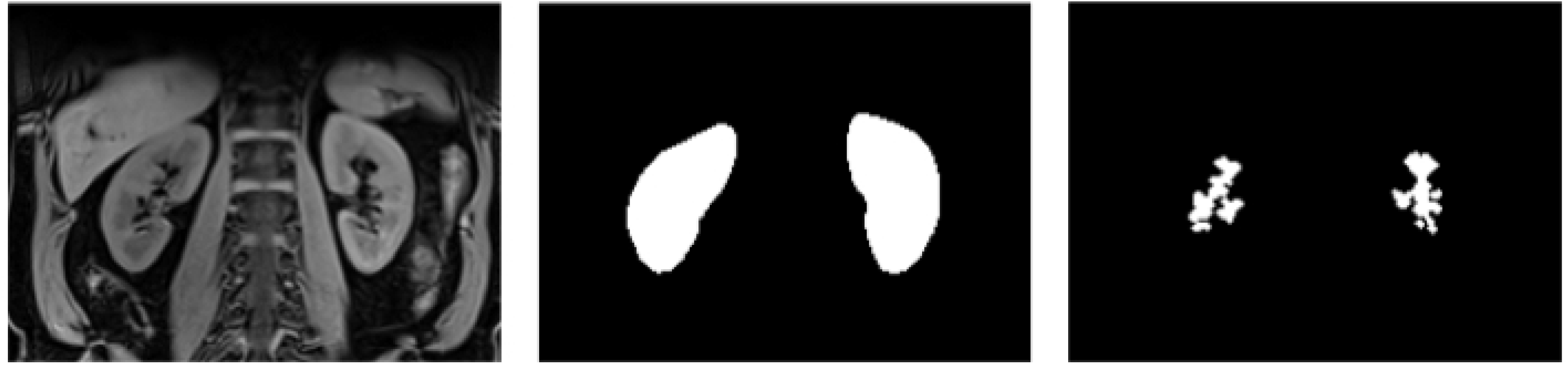
Exemplary segmentation of a coronal T1w Dixon-VIBE-dataset. (A). A whole kidney mask is generated using thresholding and active contours (B). Next the renal sinus fat is segmented using thresholding of fat isointense voxels (C).

### Covariate assessment

Covariates were obtained by standardized interviews conducted by trained staff and standardized, established laboratory methods. Briefly, glycemic status was determined by an Oral Glucose Tolerance Test (OGTT) and classified as normoglycemic control, prediabetes or diabetes according to WHO criteria: Subjects with prediabetes had impaired glucose tolerance, as defined by a normal fasting glucose concentration and a 2-h serum glucose concentration, as determined by OGTT, ranging between 140 and 200 mg/dl; and/or an impaired fasting glucose concentration, as defined by fasting glucose levels between 110 and 125 mg/dl, and a normal 2-h serum glucose concentration. Individuals with newly diagnosed diabetes were determined by a 2-h serum glucose concentration as determined by OGTT that was >200 mg/dl and/or a fasting glucose level that was 125 mg/dl. Normal controls were subjects with normal glucose metabolism with a 2-h serum glucose concentration measured by OGTT that was <140 mg/dl and a fasting glucose level that was <110 mg/dl (14). Cholesterol values were obtained from enzymatic, colorimetric assays, and albumin was measured by an immunonephelometric assay (21). GFR was calculated from serum creatinine according to the CKD-EPI definition (22), stratified by sex. Hypertension was defined as systolic blood pressure >= 140 mmHg or diastolic blood pressure >= 90 mmHg or intake of antihypertensive medication while being aware of having hypertension.

### Statistical Analysis

Demographic data, covariates and MRI-derived renal volumes, VAT and intrahepatic fat are presented as arithmetic means with standard deviation for continuous variables and counts with percentages for categorical variables. Global differences according to glycemic status (normoglycemic, prediabetes, diabetes) were determined by one-way ANOVA or χ^2^-Test, where appropriate.

Associations between glycemic status and renal volumes were determined by linear regression adjusted for additional potential confounding covariates. Regression coefficients β with corresponding 95% Confidence Intervals (CI) and p-values are reported. Furthermore, correlations between renal volumes and VAT were explored graphically by scatterplots and calculated quantitatively by Pearson’s correlation coefficient. Associations were determined by linear regression models; the corresponding R^2^ served as a measure of how much variance in the outcome was explained by the model.

All calculations were conducted with R v3.3.1. P-values < 0.05 are considered to denote statistical significance.

## Results

### Study subjects

Detailed information is available in **Table 1**. Among the 400 participants of the KORA-MRI Study, 366 (92%) subjects were included. Twenty-two (5.5%) subjects met the exclusion criteria due to none assessable datasets (9.3%), incomplete fat / water images (2%) or inadequate image quality (1%). Of the included subjects, 49 (313.4%) had diabetes and 24 prediabetes while 230 (66.7%) were normoglycemic. (**Fig 2**).

**Table 1.** Demographics, Cardiovascular Risk Factors and MRI Parameters of the Study Participants.

|  | All | Control | Prediabetes | Diabetes | P-value |
| --- | --- | --- | --- | --- | --- |
|  | N = 366 | N = 230<br>(62.8%) | N = 87<br>(23.8%) | N = 49<br>(13.4%) |  |
| Age, years | 56.2 ± 9.1 | 54.4 ± 8.9 | 58.1 ± 8.6 | 61.4 ± 8.3 | <0.001 |
| Male, % | 208 (56.8%) | 117 (50.9%) | 55 (63.2%) | 36 (73.5%) | 0.006 |
| Weight, kg | 82.9 ± 16.8 | 78.2 ± 15.5 | 91.5 ± 14.9 | 89.5 ± 17.9 | <0.001 |
| Height, cm | 171.5 ± 9.8 | 171.2 ± 10.3 | 172.3 ± 9.2 | 171.3 ± 7.9 | 0.636 |
| BMI, kg/m<sup>2</sup> | 28.1 ± 5.0 | 26.6 ± 4.3 | 30.9 ± 5.0 | 30.4 ± 5.2 | <0.001 |
| Waist circumference, cm | 98.5 ± 14.6 | 93.5 ± 12.8 | 106.6 ± 13.0 | 107.8 ± 14.5 | <0.001 |
| Waist-To-Hip-Ratio | 0.9 ± 0.1 | 0.9 ± 0.1 | 1.0 ± 0.1 | 1.0 ± 0.1 | <0.001 |
| Systolic Blood Pressure, mmHg | 120.8 ± 16.9 | 116.6 ± 15.2 | 125.8 ± 15.1 | 131.9 ± 20.4 | <0.001 |
| Diastolic Blood pressure, mmHg | 75.3 ± 10.1 | 73.6 ± 9.2 | 78.4 ± 9.4 | 78.1 ± 13.1 | <0.001 |
| Hypertension | 124 (33.9%) | 48 (20.9%) | 41 (47.1%) | 35 (71.4%) | <0.001 |
| Antihypertensive medication | 91 (24.9%) | 38 (16.5%) | 29 (33.3%) | 24 (49.0%) | <0.001 |
| Smoking |  |  |  |  | 0.339 |
| Never-smoker | 134 (36.6%) | 90 (39.1%) | 27 (31.0%) | 17 (34.7%) |  |
| Ex-smoker | 160 (43.7%) | 91 (39.6%) | 45 (51.7%) | 24 (49.0%) |  |
| Smoker | 72 (19.7%) | 49 (21.3%) | 15 (17.2%) | 8 (16.3%) |  |
| Total Cholesterol, mg/dl | 218.2 ± 37.1 | 216.1 ± 36.2 | 225.5 ± 32.0 | 214.9 ± 47.3 | 0.107 |
| HDL Cholesterol, mg/dl | 61.8 ± 18.0 | 65.4 ± 18.1 | 57.6 ± 14.5 | 51.9 ± 18.4 | <0.001 |
| LDL Cholesterol, mg/dl | 140.3 ± 33.4 | 138.5 ± 32.2 | 147.6 ± 30.1 | 136.1 ± 42.1 | 0.060 |
| Triglycerides, mg/dl | 131.5 ± 86.6 | 106.9 ± 64.3 | 153.6 ± 83.4 | 207.9 ± 123.2 | <0.001 |
| GFR (CKD-EPI) | 92.9 ± 13.0 | 94.9 ± 12.5 | 89.3 ± 12.2 | 89.6 ± 14.8 | <0.001 |
| Urine Albumin, mg/l | 6.7 [3.5, 13.0] | 6.0 [3.1, 10.6] | 6.6 [3.9, 13.8] | 11.7 [6.7, 37.9] | <0.001 |
| Visceral Fat, l | 4.5 ± 2.7 | 3.5 ± 2.3 | 5.8 ± 2.3 | 6.8 ± 2.4 | <0.001 |
| Hepatic fat (PDFF), % | 8.6 ± 7.7 | 5.6 ± 5.1 | 12.3 ± 7.8 | 16.1 ± 9.3 | <0.001 |
| Renal Volume, ml | 291.3 ± 68.7 | 280.3 ± 64.7 | 303.7 ± 67.4 | 320.6 ± 77.7 | <0.001 |
| Sinus Volume, ml | 40.0 ± 18.0 | 34.6 ± 16.0 | 47.6 ± 16.2 | 52.0 ± 19.4 | <0.001 |
| Sinus Fat, ml | 26.2 ± 13.6 | 22.2 ± 12.4 | 32.0 ± 12.0 | 34.5 ± 14.1 | <0.001 |

HDL, high-density lipoprotein; LDL, low-density lipoprotein; GFR, glomerular filtration rate, PDFF, proton density fat fraction;

**Figure 2.**
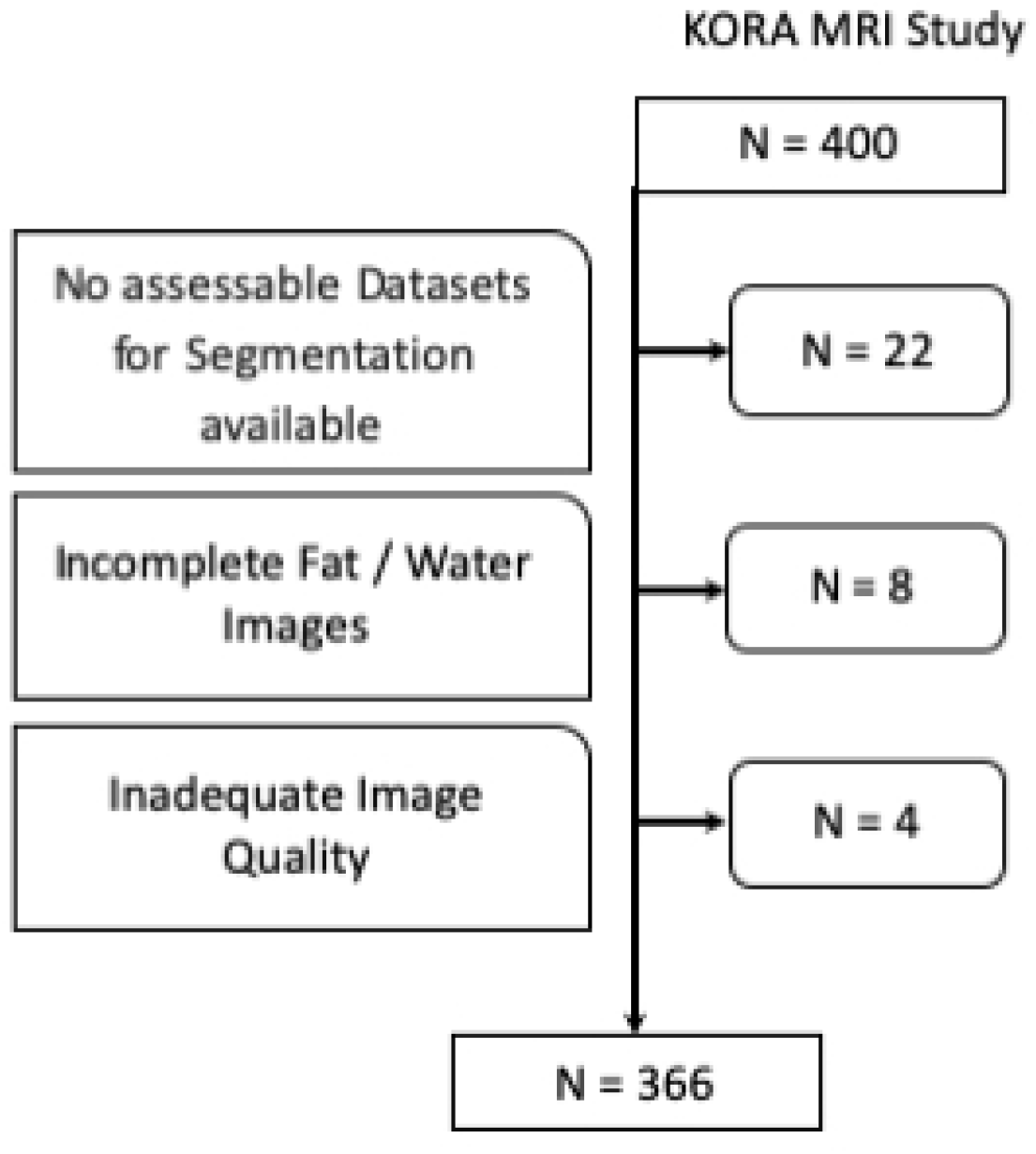
Inclusion flow chart. N = 366 study subjects were finally included for analysis.

Subjects with prediabetes and diabetes had increasing cardiovascular risk factors and metabolic syndrome components waist circumference, weight, BMI, blood pressure, blood lipids, pericardial fat and VAT. GFR showed a slight but significant decline between groups. There was no significant difference for blood albumin.

### Inter-Reader-Variability

Inter-Reader Variability was evaluated on 33 subjects. The relative difference between readers for absolute renal volume was −1.9 ml (corresponds to −0.5%, 95% limits of agreement: −29.1 ml, 25.2 ml), whereas the relative difference for renal sinus volume was −5.1 ml (corresponds to −15.2%, 95% limits of agreement: −23.3 ml, 13.0 ml) and for the percentage of renal sinus fat 7.1% (corresponds to 14.7%, 95% limits of agreement: −8.3%, 22.5%).

### Unadjusted Renal Volumes

Detailed information is provided in **Table 1** and **Fig 3**. Average renal volume showed a slight but highly significant increase between normoglycemic individuals and subjects with prediabetes and diabetes. Also, the renal sinus showed a significant enlargement between normoglycemic individuals and subjects with prediabetes and diabetes (renal volume: 280.3±64.7 ml vs 303.7±67.4 ml vs 320.6±77.7ml, respectively, p < 0.001). The largest difference was found between normoglycemic subjects and subjects with prediabetes (p<0.001 respectively). The sinus fat component showed very similar changes.

**Figure 3.**
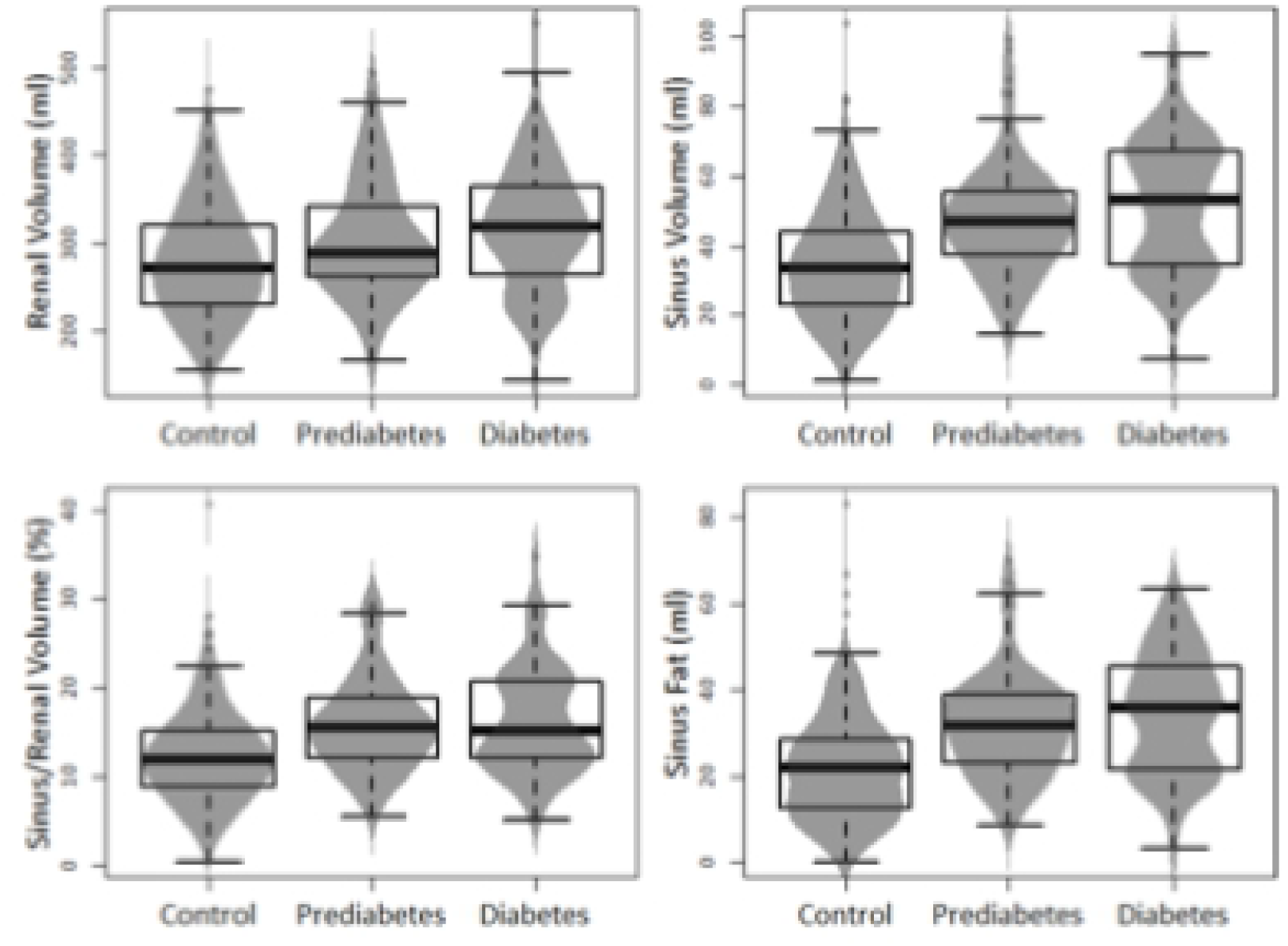
Boxplots with density curves displaying the distribution of renal and sinus fat volumes according to glycemic status. There was a considerable increase between controls and subjects with prediabetes particularly for renal sinus fat.

### Adjustment for age and sex

After adjustment for age and sex, a significant association could be found for prediabetes and diabetes with renal volume (**Table 2**). A prediabetic status was significantly associated with an increased sinus volume (β = 10.08, 95% CI: [6.5, 13.7]; p<0.01) and fat component (β = 7.13, 95% CI: [4.5, 9.8]; p<0.001). Diabetes was also significantly associated with increased sinus volume (β = 11.86, 95% CI: [7.2, 16.5]; p<0.01) and sinus fat (β = 7.34, 95% CI: [4.0, 10.7]; p<0.001). Increasing age was significantly associated with decreasing kidney volume (β = −1.35., 95% CI: [-2.0, 0.7]; p<0.01).

**Table 2.**
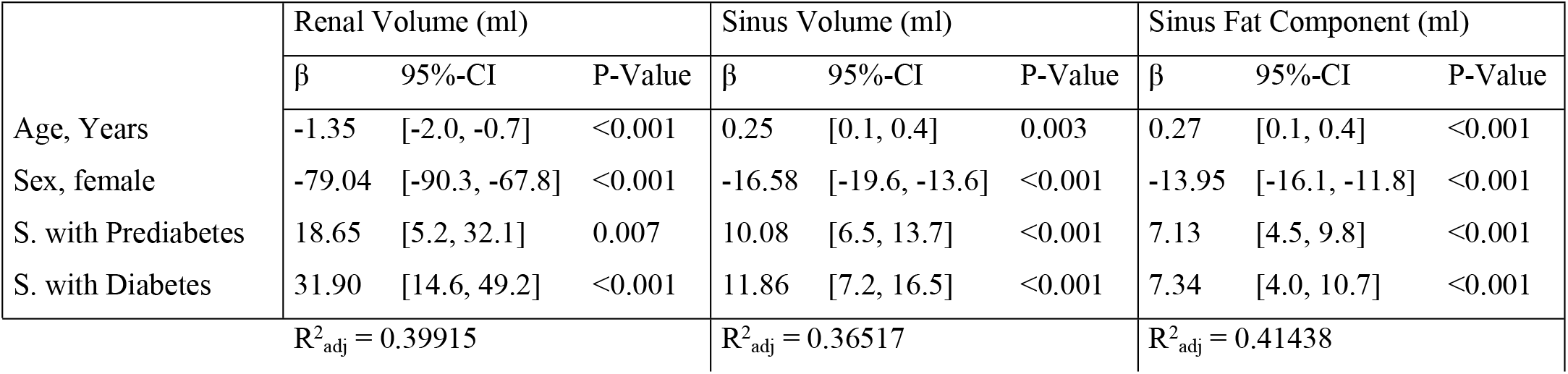
Regression model with adjustments for age, gender and glycemic status

### Adjustment for variables associated with metabolic syndrome

After adjustment for VAT, HDL, LDL, albumin, liver fat, GFR and hypertension the association between glycemic status and renal volumes decreased and was only significant for prediabetes and sinus volume (**Table 3**) (β=4.0 95% CI [0.4, 7.6]; p<0.05). Hypertension was significantly associated with increased sinus volume (β = 3.7, 95% CI: [0.4, 6.9; p<0.05]) and absolute sinus fat volume (β = 3.0, 95%CI: [0.7, 5.2]; p<0.05). GFR and all renal volumes were significantly associated as well as urine albumin levels and renal sinus volume (β = 1.6, 95% CI: [0.2, 3.0]; p<0.05).

**Table 3.** Regression model with adjustments for age, VAT, HDL, LDL, albumin, liver fat, GFR, gender, hypertension yes/no and glycemic status

|  | Renal Volume (ml) |  |  | Sinus Volume (ml) |  |  | Sinus Fat Component (ml) |  |  |
| --- | --- | --- | --- | --- | --- | --- | --- | --- | --- |
| | $\beta$ | 95%-CI | P-Value | $\beta$ | 95%-CI | P-Value | $\beta$ | 95%-CI | P-Value |
| Age, Years | 0.7 | [-6.3, 7.7] | 0.840 | 2.6 | [0.7, 4.4] | 0.006 | 2.3 | [1.1, 3.6] | <0.001 |
| VAT, l | 10.1 | [1.6, 18.7] | 0.021 | 7.5 | [5.3, 9.7] | <0.001 | 6.0 | [4.5, 7.5] | <0.001 |
| HDL | -10.5 | [-16.8, -4.2] | 0.001 | 0.2 | [-1.4, 1.8] | 0.799 | 0 | [-1.2, 1.1] | 0.948 |
| LDL | -7.6 | [-13.0, -2.2] | 0.006 | -1.3 | [-2.7, 0.1] | 0.061 | -0.5 | [-1.5, 0.5] | 0.329 |
| Urine Albumin | 2.2 | [-3.1, 7.6] | 0.414 | 1.6 | [0.2, 3.0] | 0.025 | 0.5 | [-0.5, 1.4] | 0.342 |
| Liver Fat, % | 0.2 | [-7.2, 7.5] | 0.967 | 1.6 | [-0.3, 3.5] | 0.109 | 1.6 | [0.3, 2.9] | 0.020 |
| GFR | 23.1 | [16.4, 29.7] | <0.001 | 3.0 | [1.3, 4.7] | <0.001 | 2.1 | [0.9, 3.3] | <0.001 |
| Sex, female | -57.6 | [-70.6, -44.7] | <0.001 | -8.8 | [-12.1, -5.5] | <0.001 | -7.3 | [-9.7, -5.0] | <0.001 |
| Hypertonia, yes | 10.1 | [-2.6, 22.8] | 0.118 | 3.7 | [0.4, 6.9] | 0.027 | 3.0 | [0.7, 5.2] | 0.012 |
| S. w. Prediabetes | 11.3 | [-2.6, 25.2] | 0.110 | 4.0 | [0.4, 7.6] | 0.028 | 1.7 | [-0.8, 4.3] | 0.172 |
| S. w. Diabetes | 7.7 | [-11.1, 26.5] | 0.421 | 0.0 | [-4.8, 4.8] | 1.000 | -1.7 | [-5.1, 1.7] | 0.324 |
| | $R^2_{adj} = 0.49993$ | | | $R^2_{adj} = 0.52392$ | | | $R^2_{adj} = 0.59009$ | | |

VAT, visceral adipose tissue; HDL, high-density lipoprotein; LDL, low-density lipoprotein; GFR, glomerular filtration rate, PDFF, proton density fat fraction;

### Association and Correlation between renal volumes and VAT

There was a highly significant association between VAT and renal volumes, particularly between VAT and the absolute sinus fat volume (β = 2.75, 95% CI: [2.3, 3.2]; p<0.01) (**Table 3**). A regression model only adjusted for age, sex and age already accounts for 55.6% of the variability of sinus fat (**Table 4**). When stratifying according to glycemic status, there was also a significant correlation between VAT and sinus fat in normoglycemic individuals and individuals with diabetes (between r=0.66 and 0.73) and a lower but still significant correlation in individuals with prediabetes (**Table 5** and **Fig. 4**) (between r=0.35 and 0.40)

**Table 4.** Regression model with adjustments for age, gender and VAT

|  | Renal Volume (ml) |  |  | Sinus Volume (ml) |  |  | Sinus Fat Component (ml) |  |  |
| --- | --- | --- | --- | --- | --- | --- | --- | --- | --- |
| | $\beta$ | 95%-CI | P-Value | $\beta$ | 95%-CI | P-Value | $\beta$ | 95%-CI | P-Value |
| Age, Years | -1.53 | [-2.2, -0.9] | <0.001 | 0.13 | [-0.0, 0.3] | 0.096 | 0.15 | [0.0, 0.3] | 0.006 |
| VAT | 6.06 | [3.6, 8.6] | <0.001 | 3.57 | [2.9, 4.2] | <0.001 | 2.75 | [2.3, 3.2] | <0.001 |
| Sex, female | -66.35 | [-79.4, -53.3] | <0.001 | -8.3 | [-11.5, -5.1] | <0.001 | -7.43 | [-9.7, -5.2] | <0.001 |
| | $R^2_{adj} = 0.41463$ | | | $R^2_{adj} = 0.48$ | | | $R^2_{adj} = 0.55612$ | | |

VAT, visceral adipose tissue

**Table 5.** Pearson’s correlation coefficients of VAT and renal volumes with corresponding 95% CI stratified by glycemic status

|  | Renal Volume, ml | Sinus Volume, ml | Sinus Fat Component, ml |
| --- | --- | --- | --- |
| Control | 0.42 [0.30, 0.52] | 0.67 [0.59, 0.73] | 0.73 [0.66, 0.78] |
| Patients with Prediabetes | 0.28 [0.07, 0.46] | 0.35 [0.15, 0.52] | 0.40 [0.21, 0.57] |
| Patients with Diabetes | 0.43 [0.17, 0.63] | 0.66 [0.47, 0.80] | 0.69 [0.51, 0.82] |

**Figure 4.**
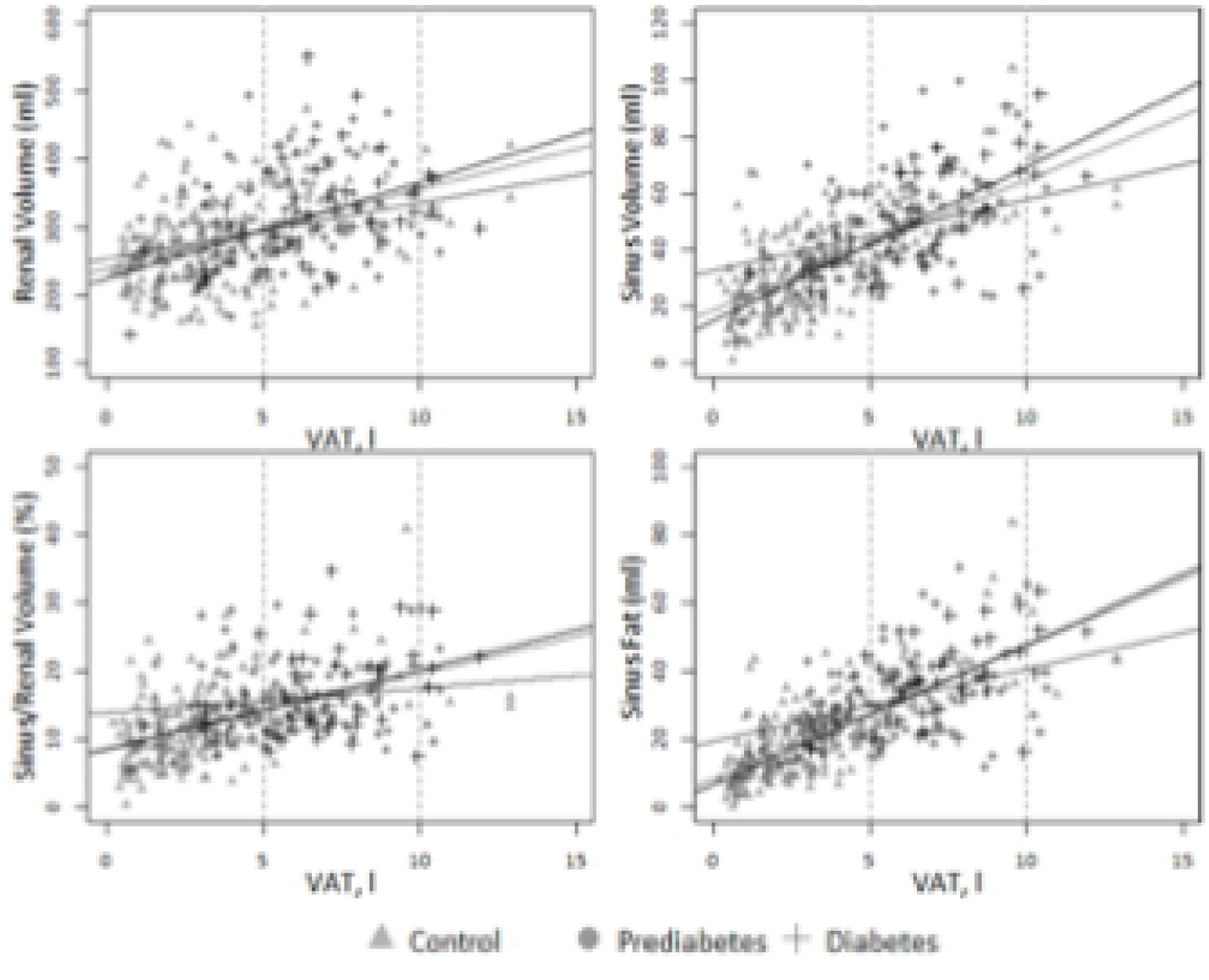
Scatter diagrams showing the correlation of the VAT with the glycemic groups. There was a significant correlation between VAT and renal sinus fat particularly for healthy controls and individuals with diabetes.

## Discussion

In our study, total renal compartment volumes significantly increased with glucose intolerance. Particularly renal sinus fat shows a considerable and significant increase in subjects with prediabetes compared to healthy controls. However, renal sinus fat is not independently associated with glycemic status and shows a strong correlation with VAT.

Assessment of kidney size and volume in the context of chronic kidney disease associated with cardiovascular risk profiles has been of long-standing interest with contradictory results (23–27). In diabetic nephropathy pre-clinical studies show an increase of kidney volume even preceding hyperfiltrative stages (28–30) similarly to our study, although this could not be verified in a small case study (31). In renovascular disease, kidney and cortex volume had a predictive value on clinical outcome (25, 26). A recent study has shown a negative correlation of the kidney volume to the extent of chronic disease (32). However, in our study cohort GFR was only slightly reduced in our well-adjusted study subjects with prediabetes and diabetes, so that extent of chronic kidney disease was only low.

Renal sinus fat has become of increasing interest when studying cardiovascular risk factors in metabolic syndrome, as perivascular adipose tissue forms an important link between obesity, insulin resistance and both macro- and microangiopathy. It is thought to obstruct lymph and blood outflow of the kidney. A recent study has shown that in a metabolically benign condition renal sinus fat reduces the release of (pro)-inflammatory factors. However, in a metabolically malignant condition, when fatty liver-derived hepatokines, like Fetuin-A, act on human renal sinus fat, the beneficial influence on glomerular cells is abolished, possibly leading to renal dysfunction and damage (33).

Renal sinus fat volume is associated with the number of prescribed antihypertensive medications and stage II hypertension (9) and is also thought to be an independent risk indicator of coronary artery calcification in middle-aged patients (34). Our results corroborate with these findings, showing that individuals with prediabetes already show a considerable increase in renal sinus fat, whereas GFR remained almost constant. Renal sinus fat may also play an early role in the pathogenesis of exercise-induced albuminuria independently of sex, age, VAT and mean arterial peak pressure (35), so that it may serve as an early imaging biomarker for potential renal disease.

Previous studies have demonstrated the presence of interindividually varying amounts of fat around the vessels of the renal hilum in humans (7, 35) potentially influencing renal/glomerular function via organ crosstalk. Accordingly, in a large subcohort of the Framingham study, quantification of renal sinus fat accumulation was independently associated with both hypertension and chronic kidney disease (7). In our study there was a significant association between increasing urine albumin levels and renal sinus volume, but not renal sinus fat volume, so that we cannot provide a final conclusion to this point.

A cross-sectional study (36) found deposition of adipose tissue particularly into the left renal sinus, which was related with the VAT amount. However, reductions in VAT volume were not accompanied by reductions in renal sinus fat accumulation. An increasing number of studies suggest, that renal sinus fat plays an important role in obesity-induced renal injury (37) which could be diagnosed and linked with early biomarkers of kidney injury (36).

In our study the volume of renal sinus fat was not independently associated with glycemic status and this association was not significant when corrected for cardiovascular risk factors. Interestingly, there was a strong correlation with VAT, explaining the major variability of renal sinus fat. These findings show, that both VAT and renal sinus fat may show interactions as perivascular adipose tissue, similarly to pericardial and hepatic fat. These findings are similar to another study investigating the same study cohort, showing that pancreatic fat content differs significantly between subjects with prediabetes, diabetes and controls, but that association is confounded by age, gender, and the amount of VAT (15).

Our data was based on semi-automated segmentation and volumetry of T1w-Dixon images. We chose a semi-automated approach as manual segmentation of abdominal organs is complex and tedious and is also prone to inter- and intraindividual bias (38). Semi-automated segmentation and volumetry of the entire kidneys may form a robust method to assess discrete changes of organ volume, which may be overlooked by a manual approach as performed in previous studies (39, 40). Our exploited algorithm was based on thresholding and geometrical approaches and did not comprise neural networks and deep learning approaches, so that manual correction was still required. However, total renal volume did only show a small inter-reader variability, whereas there was a larger relative variability for renal sinus fat, but still considerably smaller than the difference between healthy and prediabetic subjects.

There are several limitations to our study. First, our semi-automated algorithm did not satisfactory separate renal cortex and medulla (data not shown). As our focus lay on the assessment total renal volume and renal sinus fat, we did not further pursue corticomedullary discrimination and segmentation. Second, an animal study has shown, that an increase in intrarenal lipids up to 13 percent could be detected in diabetic nephropathy and associated with renal hypoxia (41). The exploited two-point-Dixon-VIBE-sequence would principally allow for such intrarenal lipid quantification, however these results are only consistent, if fat content is above ten percent, so that variation bias was too high (data not shown). More precise multi-echo-Dixon-VIBE-sequences centered on the liver were acquired in this study cohort, so that the kidneys were only partially covered. Lastly, our algorithm required manual correction, so that further refinement would be necessary to assess large volume cohort studies such as the German National Cohort (42) or UK Biobank (16) with up to 100,000 study subjects.

In conclusion, renal volume and particularly renal sinus fat volume already increases significantly in prediabetic subjects. There is a significant association between VAT and renal sinus fat, suggesting that there are metabolic interactions between these perivascular fat compartments.

## References

1. Gregory DD, Sarnak MJ, Konstam MA, Pereira B, Salem D. Impact of chronic kidney disease and anemia on hospitalization expense in patients with left ventricular dysfunction. Am J Cardiol. 2003;92(11):1300–5.

2. Sarnak MJ. Cardiovascular complications in chronic kidney disease. Am J Kidney Dis. 2003;41(5 Suppl):11–7.

3. Sarnak MJ, Levey AS, Schoolwerth AC, Coresh J, Culleton B, Hamm LL, et al. Kidney disease as a risk factor for development of cardiovascular disease: a statement from the American Heart Association Councils on Kidney in Cardiovascular Disease, High Blood Pressure Research, Clinical Cardiology, and Epidemiology and Prevention. Hypertension. 2003;42(5):1050–65.

4. Drey N, Roderick P, Mullee M, Rogerson M. A population-based study of the incidence and outcomes of diagnosed chronic kidney disease. Am J Kidney Dis. 2003;42(4):677–84.

5. Muntner P, He J, Hamm L, Loria C, Whelton PK. Renal insufficiency and subsequent death resulting from cardiovascular disease in the United States. J Am Soc Nephrol. 2002;13(3):745–53.

6. Shlipak MG, Heidenreich PA, Noguchi H, Chertow GM, Browner WS, McClellan MB. Association of renal insufficiency with treatment and outcomes after myocardial infarction in elderly patients. Ann Intern Med. 2002;137(7):555–62.

7. Lee JJ, Pedley A, Hoffmann U, Massaro JM, Levy D, Long MT. Visceral and Intrahepatic Fat Are Associated with Cardiometabolic Risk Factors Above Other Ectopic Fat Depots: The Framingham Heart Study. Am J Med. 2018;131(6):684–92 e12.

8. Siegel-Axel DI, Haring HU. Perivascular adipose tissue: An unique fat compartment relevant for the cardiometabolic syndrome. Rev Endocr Metab Disord. 2016;17(1):51–60.

9. Chughtai HL, Morgan TM, Rocco M, Stacey B, Brinkley TE, Ding J, et al. Renal sinus fat and poor blood pressure control in middle-aged and elderly individuals at risk for cardiovascular events. Hypertension. 2010;56(5):901–6.

10. Michaely HJ, Morelli JN, Budjan J, Riffel P, Nickel D, Kroeker R, et al. CAIPIRINHA-Dixon-TWIST (CDT)-volume-interpolated breath-hold examination (VIBE): a new technique for fast time-resolved dynamic 3-dimensional imaging of the abdomen with high spatial resolution. Invest Radiol. 2013;48(8):590–7.

11. Hueper K, Rong S, Gutberlet M, Hartung D, Mengel M, Lu X, et al. T2 relaxation time and apparent diffusion coefficient for noninvasive assessment of renal pathology after acute kidney injury in mice: comparison with histopathology. Invest Radiol. 2013;48(12):834–42.

12. Morrell GR, Zhang JL, Lee VS. Science to practice: Renal hypoxia and fat deposition in diabetic neuropathy--new insights with functional renal MR imaging. Radiology. 2013;269(3):625–6.

13. Holle R, Happich M, Lowel H, Wichmann HE, Group MKS. KORA--a research platform for population based health research. Gesundheitswesen. 2005;67 Suppl 1:S19–25.

14. Bamberg F, Hetterich H, Rospleszcz S, Lorbeer R, Auweter SD, Schlett CL, et al. Subclinical Disease Burden as Assessed by Whole-Body MRI in Subjects With Prediabetes, Subjects With Diabetes, and Normal Control Subjects From the General Population: The KORA-MRI Study. Diabetes. 2017;66(1):158–69.

15. Heber SD, Hetterich H, Lorbeer R, Bayerl C, Machann J, Auweter S, et al. Pancreatic fat content by magnetic resonance imaging in subjects with prediabetes, diabetes, and controls from a general population without cardiovascular disease. PLoS One. 2017;12(5):e0177154.

16. Petersen SE, Matthews PM, Bamberg F, Bluemke DA, Francis JM, Friedrich MG, et al. Imaging in population science: cardiovascular magnetic resonance in 100,000 participants of UK Biobank - rationale, challenges and approaches. J Cardiovasc Magn Reson. 2013;15:46.

17. Schlett CL, Lorbeer R, Arndt C, Auweter S, Machann J, Hetterich H, et al. Association between abdominal adiposity and subclinical measures of left-ventricular remodeling in diabetics, prediabetics and normal controls without history of cardiovascular disease as measured by magnetic resonance imaging: results from the KORA-FF4 Study. Cardiovasc Diabetol. 2018;17(1):88.

18. Storz C, Hetterich H, Lorbeer R, Heber SD, Schafnitzel A, Patscheider H, et al. Myocardial tissue characterization by contrast-enhanced cardiac magnetic resonance imaging in subjects with prediabetes, diabetes, and normal controls with preserved ejection fraction from the general population. Eur Heart J Cardiovasc Imaging. 2018;19(6):701–8.

19. Storz C, Rospleszcz S, Lorbeer R, Hetterich H, Auweter SD, Sommer W, et al. Phenotypic Multiorgan Involvement of Subclinical Disease as Quantified by Magnetic Resonance Imaging in Subjects With Prediabetes, Diabetes, and Normal Glucose Tolerance. Invest Radiol. 2018;53(6):357–64.

20. Abou-El-Ghar ME, El-Diasty TA, El-Assmy AM, Refaie HF, Refaie AF, Ghoneim MA. Role of diffusion-weighted MRI in diagnosis of acute renal allograft dysfunction: a prospective preliminary study. The British journal of radiology. 2012;85(1014):e206–11.

21. Laxy M, Knoll G, Schunk M, Meisinger C, Huth C, Holle R. Quality of Diabetes Care in Germany Improved from 2000 to 2007 to 2014, but Improvements Diminished since 2007. Evidence from the Population-Based KORA Studies. PLoS One. 2016;11(10):e0164704.

22. Inker LA, Schmid CH, Tighiouart H, Eckfeldt JH, Feldman HI, Greene T, et al. Estimating glomerular filtration rate from serum creatinine and cystatin C. New England Journal of Medicine. 2012;367(1):20–9.

23. Mogensen CE, Andersen MJ. Increased kidney size and glomerular filtration rate in untreated juvenile diabetes: normalization by insulin-treatment. Diabetologia. 1975;11(3):221–4.

24. Thelwall PE, Taylor R, Marshall SM. Non-invasive investigation of kidney disease in type 1 diabetes by magnetic resonance imaging. Diabetologia. 2011;54(9):2421–9.

25. Cheung CM, Chrysochou C, Shurrab AE, Buckley DL, Cowie A, Kalra PA. Effects of renal volume and single-kidney glomerular filtration rate on renal functional outcome in atherosclerotic renal artery stenosis. Nephrol Dial Transplant. 2010;25(4):1133–40.

26. Cheung CM, Shurrab AE, Buckley DL, Hegarty J, Middleton RJ, Mamtora H, et al. MR-derived renal morphology and renal function in patients with atherosclerotic renovascular disease. Kidney Int. 2006;69(4):715–22.

27. Gandy SJ, Armoogum K, Nicholas RS, McLeay TB, Houston JG. A clinical MRI investigation of the relationship between kidney volume measurements and renal function in patients with renovascular disease. Br J Radiol. 2007;80(949):12–20.

28. Christiansen T, Rasch R, Stodkilde-Jorgensen H, Flyvbjerg A. Relationship between MRI and morphometric kidney measurements in diabetic and non-diabetic rats. Kidney Int. 1997;51(1):50–6.

29. Christiansen T, Stodkilde-Jorgensen H, Klebe JG, Flyvbjerg A. Changes in kidney volume during pregnancy in non-diabetic and diabetic rats measured by magnetic resonance imaging. Exp Nephrol. 1998;6(4):302–7.

30. Bak M, Thomsen K, Christiansen T, Flyvbjerg A. Renal enlargement precedes renal hyperfiltration in early experimental diabetes in rats. J Am Soc Nephrol. 2000;11(7):1287–92.

31. Avram MM, Hurtado H. Renal size and function in diabetic nephropathy. Nephron. 1989;52(3):259–61.

32. Woodard T, Sigurdsson S, Gotal JD, Torjesen AA, Inker LA, Aspelund T, et al. Segmental Kidney Volumes Measured by Dynamic Contrast-Enhanced Magnetic Resonance Imaging and Their Association With CKD in Older People. Am J Kidney Dis. 2014.

33. Wagner R, Machann J, Guthoff M, Nawroth PP, Nadalin S, Saleem MA, et al. The protective effect of human renal sinus fat on glomerular cells is reversed by the hepatokine fetuin-A. Sci Rep. 2017;7(1):2261.

34. Murakami Y, Nagatani Y, Takahashi M, Ikeda M, Miyazawa I, Morino K, et al. Renal sinus fat volume on computed tomography in middle-aged patients at risk for cardiovascular disease and its association with coronary artery calcification. Atherosclerosis. 2016;246:374–81.

35. Wagner R, Machann J, Lehmann R, Rittig K, Schick F, Lenhart J, et al. Exercise-induced albuminuria is associated with perivascular renal sinus fat in individuals at increased risk of type 2 diabetes. Diabetologia. 2012;55(7):2054–8.

36. Krievina G, Tretjakovs P, Skuja I, Silina V, Keisa L, Krievina D, et al. Ectopic Adipose Tissue Storage in the Left and the Right Renal Sinus is Asymmetric and Associated With Serum Kidney Injury Molecule-1 and Fibroblast Growth Factor-21 Levels Increase. EBioMedicine. 2016;13:274–83.

37. Irazabal MV, Eirin A. Role of Renal Sinus Adipose Tissue in Obesity-induced Renal Injury. EBioMedicine. 2016;13:21–2.

38. Attenberger UI, Sourbron SP, Notohamiprodjo M, Lodemann KP, Glaser CG, Reiser MF, et al. MR-based semi-automated quantification of renal functional parameters with a two-compartment model--an interobserver analysis. Eur J Radiol. 2008;65(1):59–65.

39. Will S, Martirosian P, Wurslin C, Schick F. Automated segmentation and volumetric analysis of renal cortex, medulla, and pelvis based on non-contrast-enhanced T1- and T2-weighted MR images. Magma. 2014.

40. Winter KS, Helck AD, Ingrisch M, Staehler M, Stief C, Sommer WH, et al. Dynamic Contrast-Enhanced Magnetic Resonance Imaging Assessment of Kidney Function and Renal Masses: Single Slice Versus Whole Organ/Tumor. Invest Radiol. 2014.

41. Peng XG, Bai YY, Fang F, Wang XY, Mao H, Teng GJ, et al. Renal lipids and oxygenation in diabetic mice: noninvasive quantification with MR imaging. Radiology. 2013;269(3):748–57.

42. Bamberg F, Kauczor HU, Weckbach S, Schlett CL, Forsting M, Ladd SC, et al. Whole-Body MR Imaging in the German National Cohort: Rationale, Design, and Technical Background. Radiology. 2015;277(1):206–20.

